# The covariance shift (C-SHIFT) algorithm for normalizing biological data

**DOI:** 10.1101/2020.04.13.038463

**Authors:** Evgenia Chunikhina, Paul Logan, Yevgeniy Kovchegov, Anatoly Yambartsev, Debashis Mondal, Andrey Morgun

## Abstract

Omics technologies are powerful tools for analyzing patterns in gene expression data for thousands of genes. Due to a number of systematic variations in experiments, the raw gene expression data is often obfuscated by undesirable technical noises. Various normalization techniques were designed in an attempt to remove these non-biological errors prior to any statistical analysis. One of the reasons for normalizing data is the need for recovering the covariance matrix used in gene network analysis. In this paper, we introduce a novel normalization technique, called the covariance shift (C-SHIFT) method. This normalization algorithm uses optimization techniques together with the blessing of dimensionality philosophy and energy minimization hypothesis for covariance matrix recovery under additive noise (in biology, known as the bias). Thus, it is perfectly suited for the analysis of logarithmic gene expression data. Numerical experiments on synthetic data demonstrate the method’s advantage over the classical normalization techniques. Namely, the comparison is made with rank, quantile, cyclic LOESS (locally estimated scatterplot smoothing), and MAD (median absolute deviation) normalization methods.

---

Gene expression analysis plays an important role in genomic research. Several omics technologies such as RNAseq and microarrays allow for the collection of massive amounts of simultaneous measurements of gene expression levels of thousands to tens of thousands of genes. Analyzing different patterns of gene expressions helps to gain insight into complex biological phenomena such as development, aging, onset and progression of diseases, and cellular response/reaction to drugs/treatments. Although new technologies are constantly developing, it is well known that all of them generate some technical noise which affects the measured gene expression levels [8, 20]. To extract accurate biological information it becomes necessary to normalize the data to filter out/compensate for these nonbiological noises/errors. Normalization is a crucial pre-processing step in the gene expression data analysis. The gene expression data will vary significantly after different normalization methods. Thus, the results of further data analysis (e.g. gene expression network) will be critically dependent on a choice of a normalization technique. A variety of normalization procedures have been used on gene expression data sets. See [3, 4, 12, 13, 15, 17, 19] and reference therein for a review and comparison of current normalization strategies. In this paper we develop a novel normalization technique, called the covariance shift (C-SHIFT) method, and compare it to the following well known normalization methods used in large scale data analysis: rank, quantile, cyclic LOESS (locally estimated scatterplot smoothing), and MAD (median absolute deviation). See [1, 4, 14, 15] and references therein for more details on the above listed normalization methods.

Consider a situation where the gene expression data is subjected to multiplicative noise (aka bias). Specifically, let 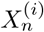 be the true gene expression, where subscript index *n* stands for the *n*-th gene in the network and the superscript index *i* stands for the *i*-th measurement. The observed gene expression, denoted by 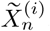, is different from 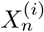 due to all gene expressions in the *i*-th measurement being distorted by i.i.d. multiplicative noise variable *W* ^(*i*)^, i.e.,

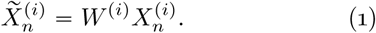

Here, both the observed and the true gene expressions are positive, i.e., 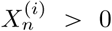 and *W*^(*i*)^ > 0. The random variables 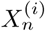 are independent of the variable *W*^(*i*)^.

In biology, the multiplicative noise *W*^(*i*)^ is referred to as the bias. The bias is prompted by random events causing an error in the measurement of the total amount of RNA. Such random events are often related to different levels of tissue preservation in different sample that leads to variability of RNA degradation. Consequently, this leads to an RNA detection problem. Additionally, there are other technical reasons for an error in the measurement of the total amount of RNA in a given sample that may lead to a bias in (1). All other noise (e.g. misreading parts of RNA) goes into the variable 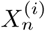

The multiplicative noise in (1) implies the corresponding additive noise (bias) in the logarithimic gene expression data:

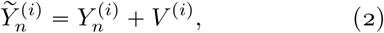

where we let 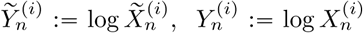, and *V*^(*i*)^ ≔ log *W*^(*i*)^.

The bias, whether multiplicative as in (1) or additive as in (2), causes the correlations to be shifted away from −1. Indeed, since

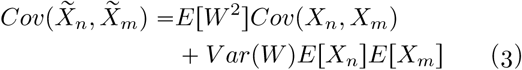

the correlation of observed gene expressions

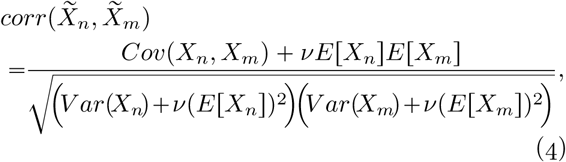

where 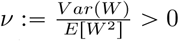. Notice that if *Cov*(*X*_*n*_, *X*_*m*_) is negative, by adding positive multiples of *v* > 0 in the numerator and the denominator as in (4), we arrive at 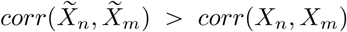. In other words, all negative correlations *corr*(*X*_*n*_, *X*_*m*_) will either turn into positive, or negative of smaller magnitude in the observed variables 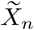. While the multiplicative bias *W*^(*i*)^ ≈ 1 (similarly, the additive bias *V*^(*i*)^ ≈ 0) may not appear critical, they are known to cause significant problems in gene correlation structure analyses. Specifically, this phenomenon is known to cause the disappearance of the large magnitude negative correlations in the observed biological data, 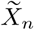, which hampers the ability to perform certain types of statistical data analysis, such as the false discovery rate (FDR) method.

The same is observed for the logarithmic data (2). Similarly to (3), the independent additive noise in (2) implies an increase of covariance,

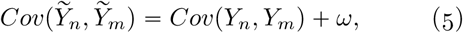

where *ω* = *Var*(*V*) > 0. Consequently, the correlations in the logarithmic data are

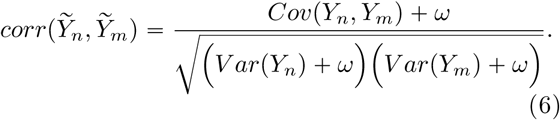

Here too, if *Cov*(*Y*_*n*_, *Y*_*m*_) is negative, by adding *ω* > 0 in the numerator and the denominator, we obtain

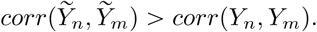

Thus, the phenomenon of disappearance of the large magnitude negative correlations also applies to the logarithmic data 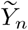.

Denote by 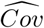 the empirical covariances taken over *N* subjects for each of 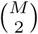 pairs of genes. Similarly, let 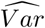 denote the empirical variance. Then, equation (2) yields the observed empirical covariance

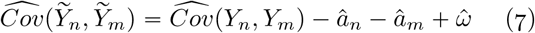

for all pairs of gene indices *n* and *m*, where 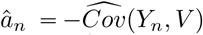 for all *n* = 1, …, *M*, and 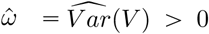. As is often the case, 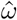 can be very large relative to the values of *â*_*n*_, causing the disappearance of the large magnitude negative correlations in empirical data.

The goal of the covariance shift (C-SHIFT) normalization method introduced here is the recovery of the true empirical covariances 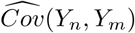 and the respective true empirical correlations in the case of the logarithmic gene expression data or any other situations with additive noise as in (2).

Let 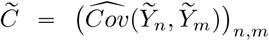 be the empirical covariance matrix of the observed data 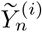, and let 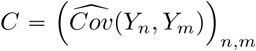 be the empirical covariance matrix of the cleaned data 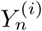 (i.e., the true empirical covariance) that we desire to recover. Formula (7) rewritten in the matrix form states

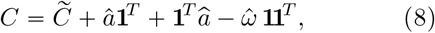

where *â* = (*â*_1_, …, *â*_*M*_)^*T*^, and **1** denotes the column vector of 1’s, hence **11**^*T*^ is a square matrix of 1’s.

Our goal here is to estimate *â* and 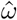 in (8), and thus recover the true empirical covariance matrix *C*. This will be done in Section I. We assume large dimension *M*. There will be two cases.

*Case I:* If 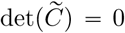 (e.g. *N* < *M*), we make a small perturbation of the diagonal entries of 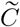 (the variances) resulting in a new covariance matrix being positive definite whose smallest eigenvalue is still very close to zero. Next, we use energy minimization to estimate *â*_*n*_ and 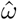 in (8).

*Case II:* If 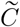 is positive definite, our approach exploits the phenomenon sometimes referred to as the *curse of dimensionality* [2, 16] and sometimes as the *blessing of dimensionality* [5, 7, 10], postulating that in higher dimensions almost all data points are located near extrema (i.e., in the outer shell)^*^. In other words, for large *M*, we anticipate the smallest eigenvalue of *C* to be near zero. As a rigorous bound, we observe that if some of the correlations *corr*(*Y*_*n*_, *Y*_*m*_) are located in [−1, *δ* − 1] interval, then the smallest eigenvalue of *C* is located within 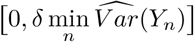 interval. Thus, as in Case I, under the blessing of dimensionality assumption, we again use energy minimization for estimating *â*_*n*_ and 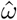.

The problem of improving the existing and developing new normalization methods is very important for scientists working with biological data. The fact that normalization alters the data-correlation structure was stated in Saccenti [19]. Besides [19] gives a comprehensive overview of normalization methods. In Bolstad *et al*. [4] the authors compare three complete data normalization methods (cyclic loess, contrast based method, and quantile) that make use of data from all arrays in an experiment to two methods that make use of a baseline array. The comparison was done on two publicly available datasets with the results favoring the complete data methods. For more on the normalization methods, see [1, 6, 8, 9, 15, 18, 21].

## I. Theoretical Derivations

### Proposition 1.

*Suppose* ℳ *is a symmetric positive definite square matrix. Then*,

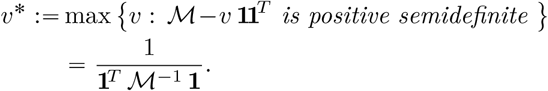

*Proof*. Observe that

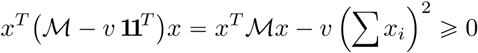

for all *x* ∈ ℝ^*M*^ if and only if *v* ⩽ *v*^*^, where *v*^*^ minimizes *x*^*T*^ ℳ*x* under the condition ∑*x*_*i*_ = *Const*. Next, applying the Lagrange multipliers method, we obtain 2ℳ*x* = λ**1**, and therefore

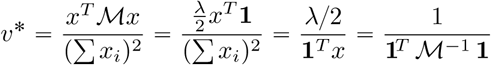

as 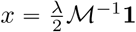. □

Suppose the empirical covariance matrix 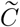 is positive definite, i.e., 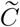 is of full rank. Consider values of a column vector *α* = (*α*_1_, …, *α*_*M*_)^*T*^ such that 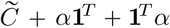 is positive definite. Then, by Prop. 1,

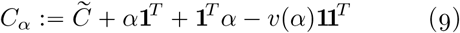

is positive semidefinite with det (*C*_*α*_) = 0, where we let

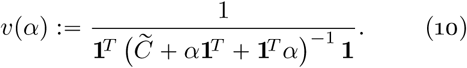

Next, recall the quantities *ã* and 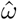 in (8). If 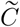 is rank deficient, we perturb its diagonal entries by adding small positive (random or deterministic) values, and if 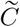 is positive definite, we assume the blessing of dimensionality phenomenon holds. Thus, in either case, we work under assumption that 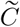 is positive definite with its smallest eigenvalue located near zero. Then, Prop. 1 implies 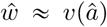, where *v*(*α*) is as defined in (10). Therefore, letting *α* = *â* in (9), we will have *C*_*â*_ approximating *C*.

Finally, for all *X* ∈ ℝ^*M*×*N*^, let ‖*X*‖_*F*_ denote the Frobenius norm of *X* and let 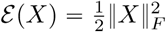 be the energy function. Our next assumption states that *â* can be estimated by the minimizer *α*^*^ of the energy function *ε*(*C*_*α*_), i.e., we estimate *â* by

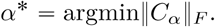

The assumption is additionally justified by the observation that a random adjustment of the covariance via an additive noise (bias) as in (7) will result in an energy increase, i.e., 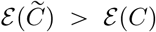. Hence, we use 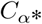 to approximate *C*_*â*_ and the desired true empirical covariance matrix *C*.

### Lemma 1.

*Suppose the empirical covariance matrix* 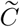 *is of full rank, and the quantities C*_*α*_, *and v(α*), *are as in* (9) *and* (10). *Then, the gradient of the Frobenius norm squared is given by*

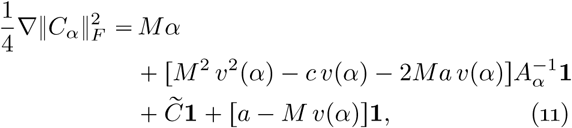

*where* ‖ · ‖ _*F*_ *denotes the Forbenius norm, and we let*

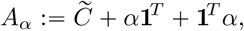

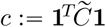 *and* 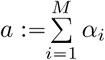.

*Proof*. By (9), we have

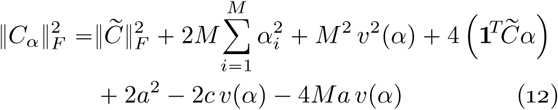

Notice that

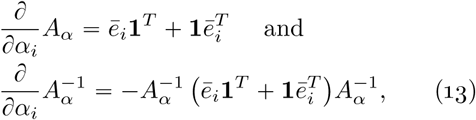

where *ē*_*i*_ is the *i*-th coordinate vector. Therefore, we have

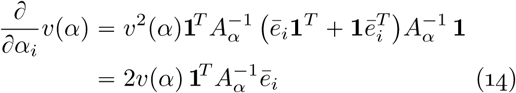

implying

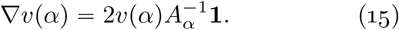

Next, the gradient 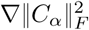 in (11) is found via the equations (12) and (15). □

First, observe that *C*_*α*_ is invariant under the addition of multiples of **1**. Thus, without loss of generality, we restrict the domain to a hyperplane *a* “Const. Next, observe that 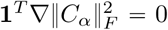 in (11). Thus, in the gradient descent method, the value of *a* remains constant, i.e., throughout the algorithm, vector *α* remains on the same hyperplane *a* “Const.

### Lemma 2.

*Suppose the empirical covariance matrix* 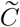 *is of full rank, and the quantities C*_*α*_, *v* (*α*), *and A*_*α*_ *are as in* (9), (10), *and* (12) *respectively. Then, the Hessian of* 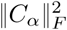, *denoted by* 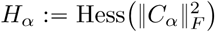 *is expressed as follows*

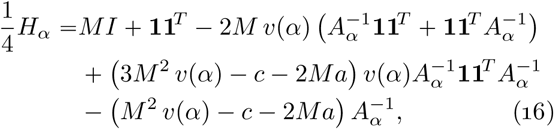

*where I is the identity matrix*, 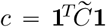, *and* 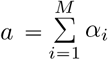.

*Proof*. By (11), we have

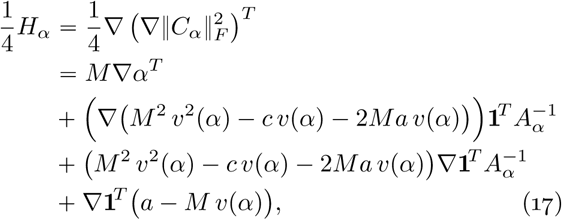

where 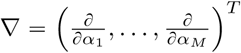 was used as the column vector of the partial derivative operators. The summation parts in (17) are calculated as follows. First,

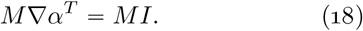

Next, (15) implies

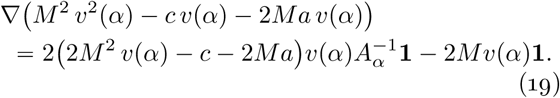

Equation (13) implies

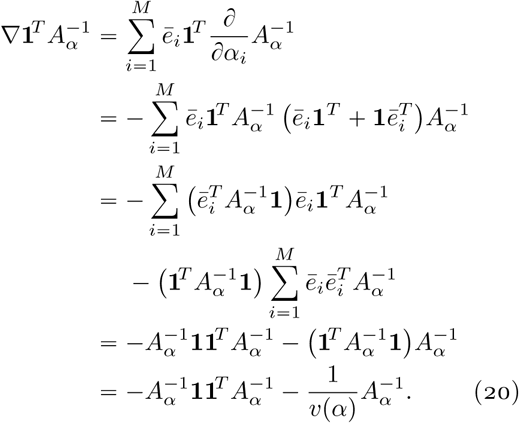

Finally, (15) is used to derive

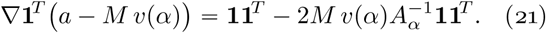

Combining together equations (18)-(21) and substituting them into (17) we obtain (16). □

### Theorem 1.

*Suppose the empirical covariance matrix* 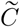 *is of full rank, and the quantities C*_*α*_ *and v(α) are as in* (9) *and* (10). *Then, the Frobenius norm squared* 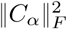 *is convex, i*.*e*.,

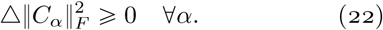

*Proof*. We will use the notations of this section such as 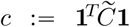 and 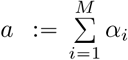. Without loss of generality we consider *α* on the hyperplane *a* = 0.

Here, 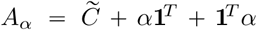 is a positive definite symmetric matrix with eigenvalues

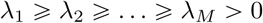

counted with respect to algebraic multiplicity, and let {*v*_*i*_}_*i* = 1,…,*M*_ be the corresponding orthonormal basis of eigenvectors.

Equation (16) implies

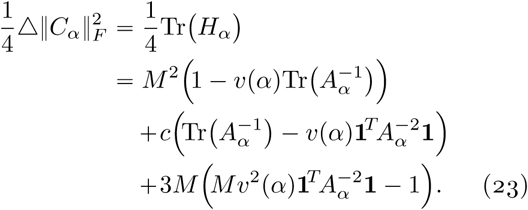

The Laplacian in (23) is shown to be strictly positive in the following three steps. First, by the Cauchy-Bunyakovsky-Schwarz inequality, we have

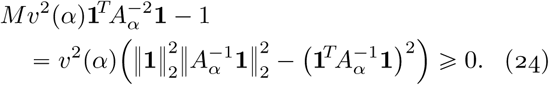

Next, observe that for *M* ⩾ 2,

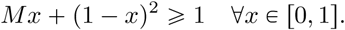

Thus, for a given probability mass function {*p*_*k*_}_*k*=1,…,*M*_ such that *p*_*k*_ < 1 for all *k*, and a given index *i* ∈ {1, …, *M*}, Jensen’s inequality implies

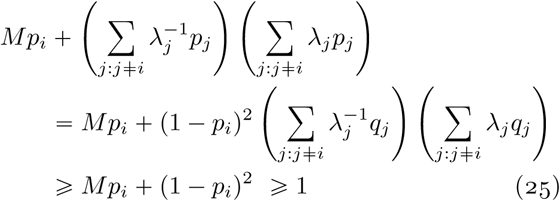

where we let 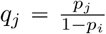 for all *j* ≠ *i*. Summing over all *i* in (25), we obtain,

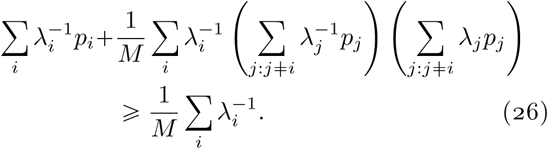

Eqn. (26) implies

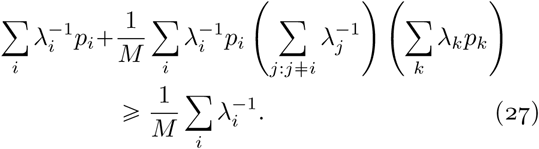

which rewrites as

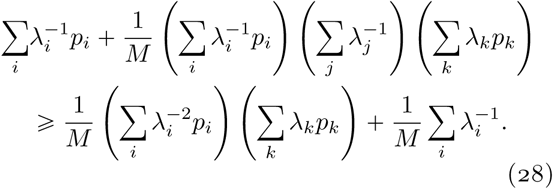

Finally, we let 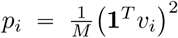 and substitute the following expressions into (28):

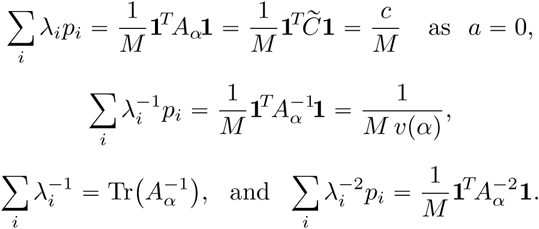

Consequently, (28) rewrites as

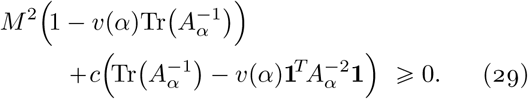

Substituting (24) and (29) into (23), we then obtain 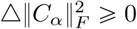. □

## II. C-SHIFT algorithm and experiments

In this section we provide the C-SHIFT algorithm and evaluate its performance on synthetic data sets. Moreover, we compare the C-SHIFT algorithm with the well-known and frequently used normalization methods: Quantile, Rank, LOESS, and Median absolute deviation (MAD). Our empirical results demonstrate that the C-SHIFT algorithm outperforms other methods.

### A. C-SHIFT algorithm

The pseudocode for the C-SHIFT algorithm is given in Algorithm 1. Note that the algorithms takes into account both cases: when 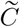 has full rank and when 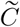 is rank deficient (i.e., 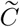 is positive semi-definite but not positive definite). When 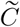 is rank deficient the rank of 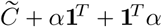 may exceed the rank 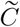 by no more than 2, and therefore may also be rank deficient. Therefore, to make 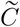 a full rank we add to it a diagonal matrix `diag`(*f*), where *f* is a vector of i.i.d. random variables from `Unif` [0, 1].

To find the optimal 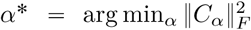, we use gradient and Hessian, provided in equations (11) and (16), in the trust-region algorithm to minimize 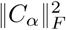.

### B. Numerical experiments

In this section we conduct experiments on two synthetic datasets that we generate using random covariance method (RCM) and cascade method. We start by describing both methods.

#### 1) Data generation

##### Algorithm 1: C-SHIFT

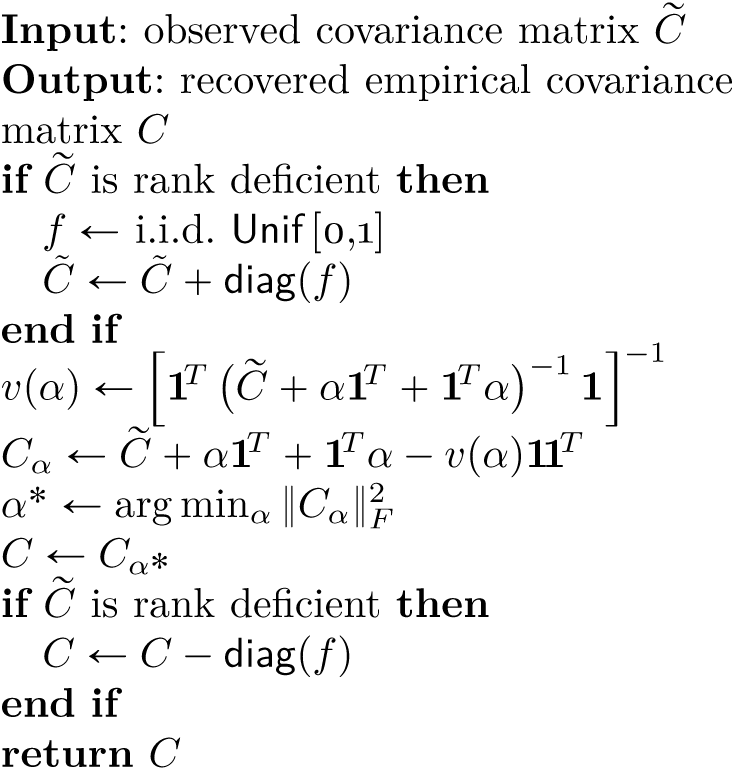

##### a) Random covariance method (RCM)

We generate a synthetic data set with *M* = 2000 genes and *N* = 50 measurements (samples) using RCM. For that we first generate an auxiliary matrix *H* ∈ ℝ^*M*×*m*^ (*m* = 2) whose entries are independent random variables, uniformly distributed over the interval *I* = [−10, 10]. Next, we sample a diagonal matrix *D* ∈ ℝ^*M*×*M*^ with diagonal entries being i.i.d. exponential random variables with parameter λ_*D*_ = 30. We let Σ = *HH*^*T*^ + *D* be the population (parameter) covariance matrix. Then we generate the true empirical logarithmic data 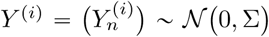 for each *i* = 1, …, *N*. Finally, we set the observed logarithmic data be 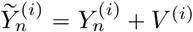, where vector *V*^(*i*)^ are *N*(− 0.01, 100) random variables.

##### b) Cascade method

The *cascade* datasets were generated according by a directed acyclic weighted network *G* = (*V, E*) aka directed acyclic graph (DAG). The graph was randomly generated via a recurrent *cascade* model. The parent-offspring relation is represented by the direction of edges *E* = {(*u, v*)} of the graph *G*, i.e., *u* is the parent vertex and *v* is its offspring. For any vertex *v* let *pa*(*v*) be the set of its parents, *pa*(*v*) = {*u* ∈ *V*: (*u, v*) ∈ *E*}. Next, for each edge (*u, v*) ∈ *E* an independent random weight *c*_*uv*_ is assigned, with c.d.f.

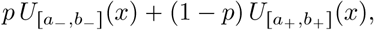

where the parameters *a*_−_ < *b*_−_ ⩽ 0, 0 ⩽ *a*_+_ < *b*_+_, and *p* ∈ (0, 1) are fixed, and *U*_*A*_(*x*) denotes the uniform c.d.f. on an interval *A*. We generated a random weighted DAG with the nodes *v* ∈ *V* representing the genes. The random variables {*Y*_*v*_}_*v*∈*V*_ representing the logarithmic gene expressions are generated as a noisy multiplicative cascade via the following structural linear recursive equations:

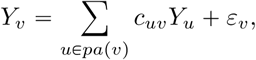

where the recursion begins with *Y*_0_ = *y*_0_, and proceeds from generation to generation. The noise variables (*ε*_*v*_, *v* ∈ *V*) are i.i.d. *N*(0, *σ*^2^), sampled independently from the random weights *c*_*uv*_. For simulation of (*Y*_*v*_, *v* ∈ *V*) the following values of parameters were chosen:

| $p$ | $[a_-, b_-]$ | $[a_+, b_+]$ | $\sigma^2$ | $y_0$ | $\vert V\vert $ |
| --- | --- | --- | --- | --- | --- |
| 1/3 | $[-1.2, -0.5]$ | $[0.5, 1.3]$ | 1 | 4.5 | 2000 |

#### 2) Simulation results

We generate two data sets (RCM and Cascade) using the methods described in section II-B1. Each date set consists of a matrix with the empirical data 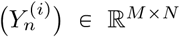 and a matrix with the observed data 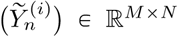. In both, RCM and Cascade data sets, we let *M* = 2000 genes and *N* = 50 measurements (samples). For each data set, we normalize the covariance matrix 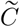, obtained from the observed data, by using C-SHIFT, Rank, Quantile, LOESS, and MAD methods. We compare the performance of the algorithms using the results presented in Figures 1-4.

**Fig. 1.**
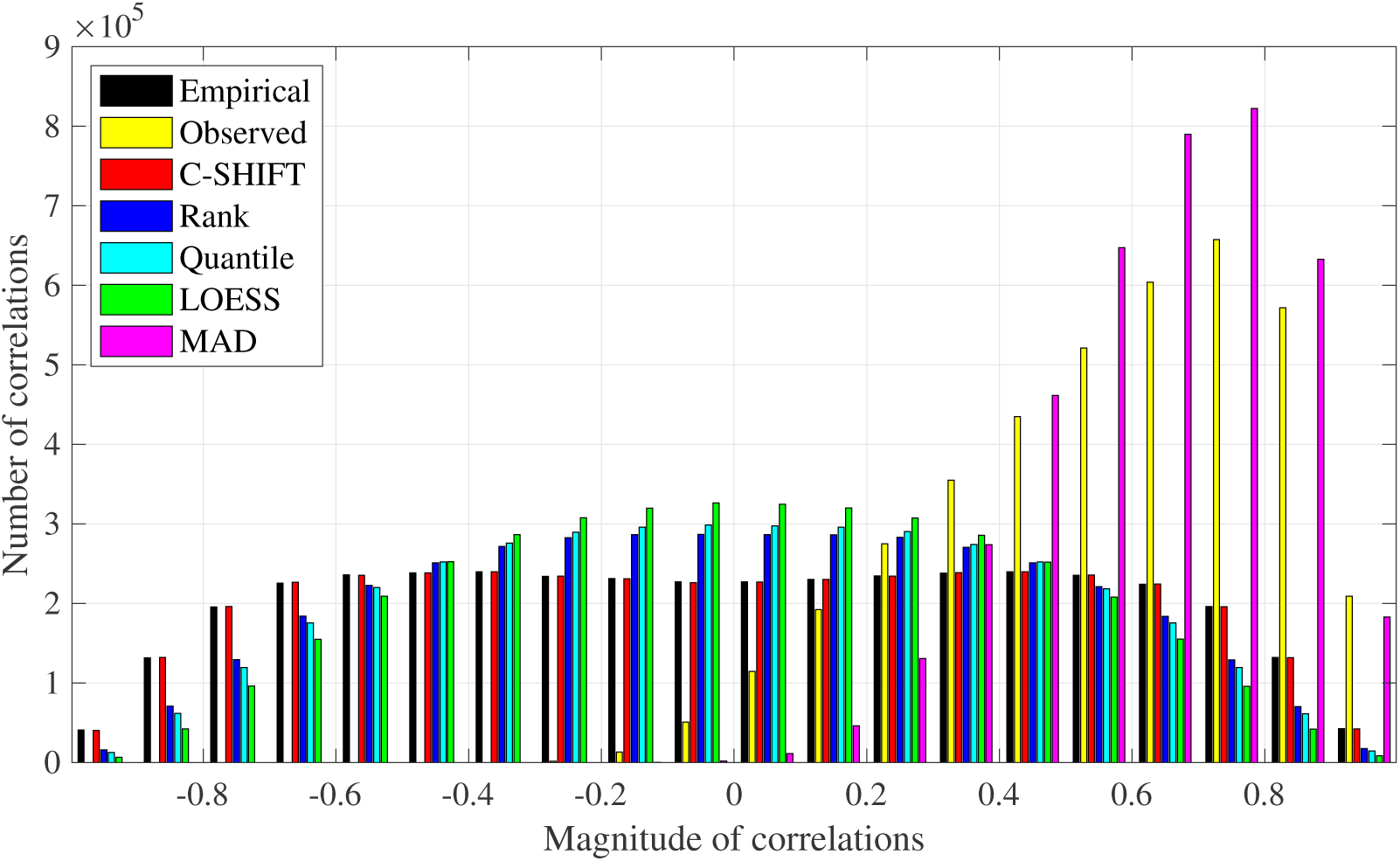
A bar graph of correlations for the RCM data set. On the x-axis we display the range of correlations, partitioned into intervals of length 0.1. The height of each bar describes the number of correlations that belong to the corresponding interval. Bars of different colors correspond to different correlation matrices, indicated in the legend.

**Fig. 2.**
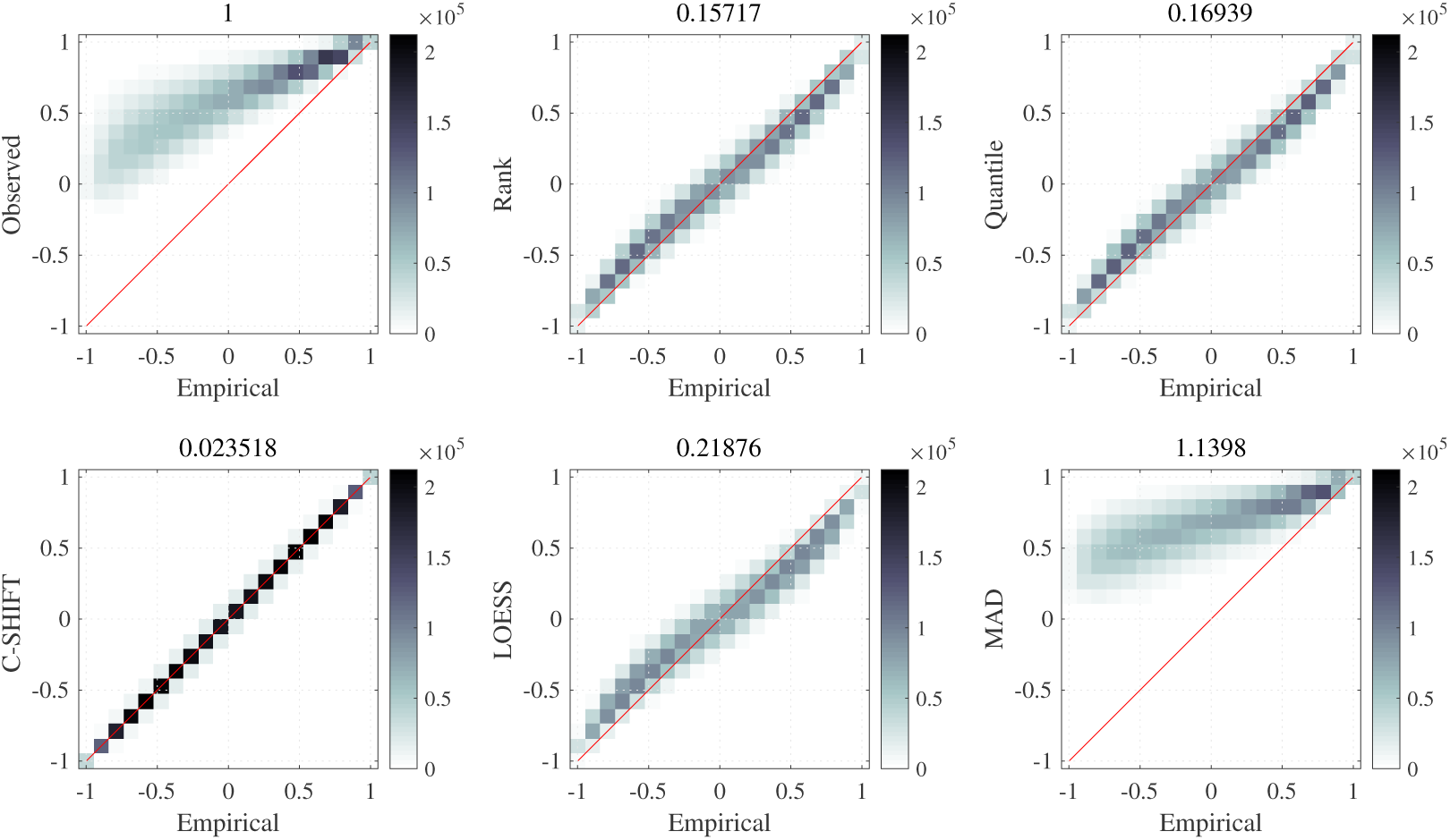
The heat maps for the RCM data set. Each heat map illustrates the transformation of the true empirical correlations *corr*(*Y*_*n*_, *Y*_*m*_) (horizontal axis) after adding bias and applying the corresponding normalization method. In the top left plot the vertical axis represents the observed correlations 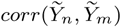. In the remaining five heat maps, the vertical coordinates represent the correlations after normalization. Going clockwise, these five heat maps are Rank, Quantile, MAD, LOESS, and C-SHIFT. The darker the color, the higher the density. The number on top of each heat map indicates the relative leftover error after normalization. Smaller numbers indicate better recovery performance.

**Fig. 3.**
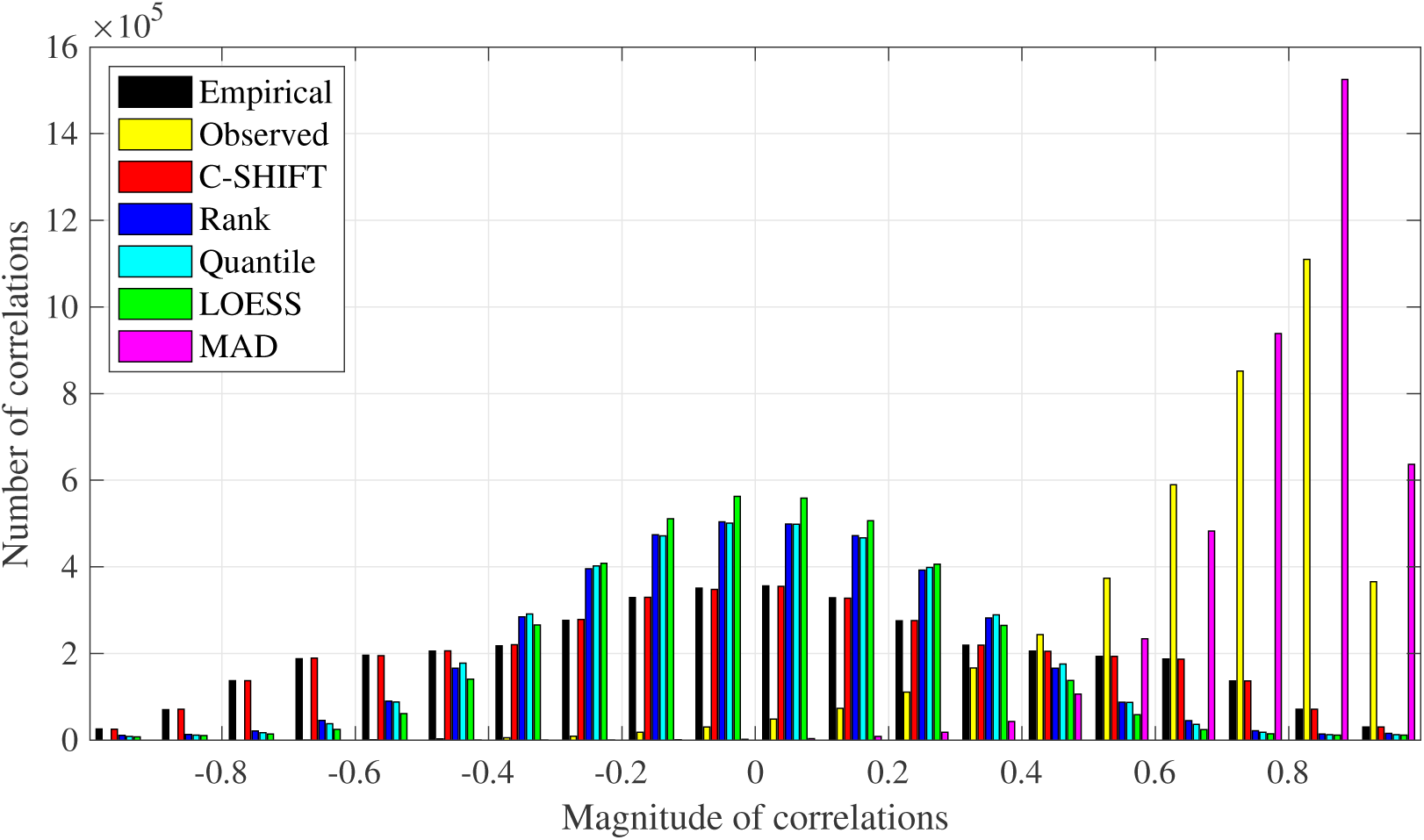
A bar graph of correlations for the Cascade data set. On the x-axis we display the range of correlations, partitioned into intervals of length 0.1. The height of each bar describes the number of correlations that belong to the corresponding interval. Bars of different colors correspond to different correlation matrices, indicated in the legend.

**Fig. 4.**
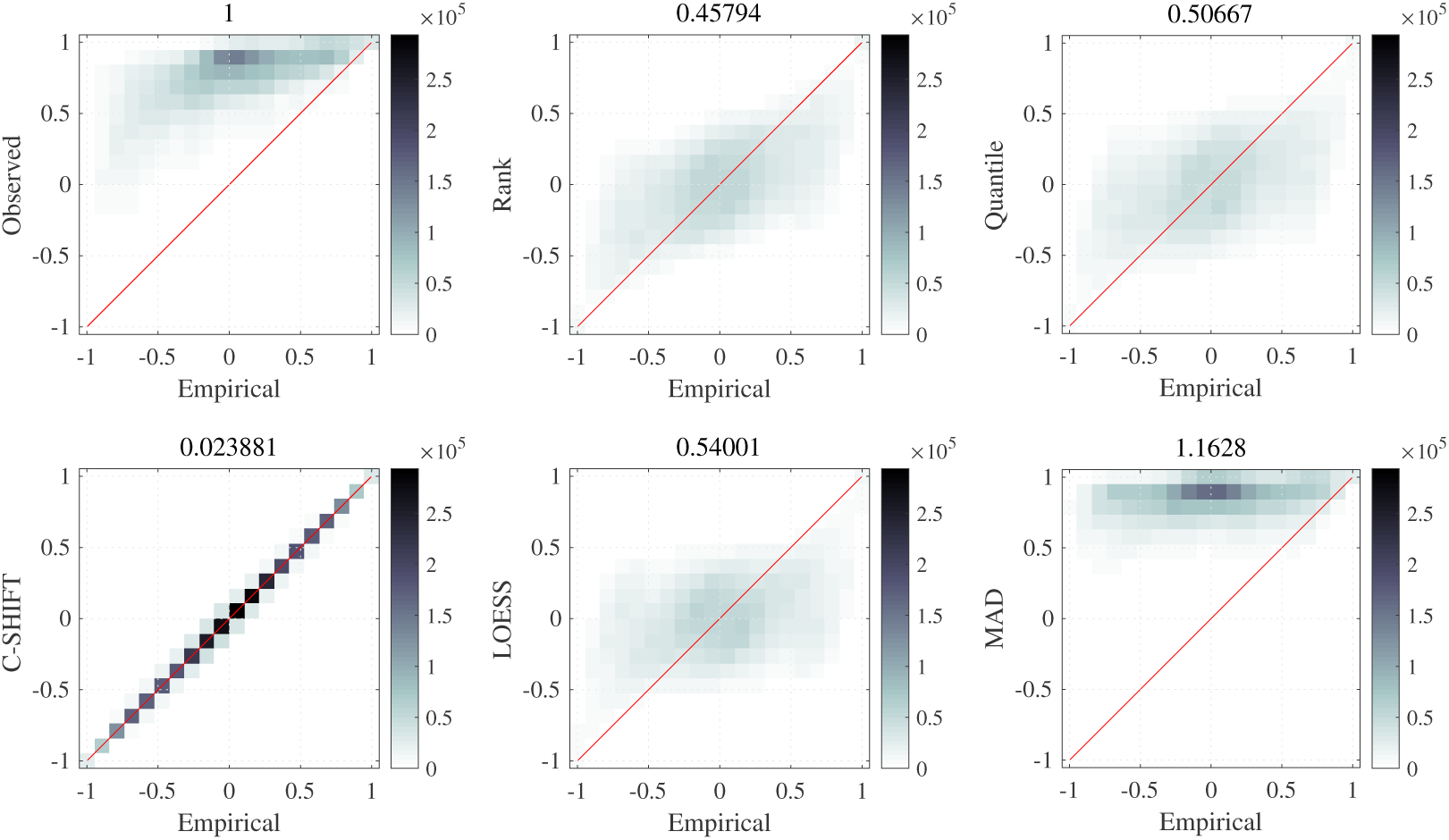
The heat maps for the Cascade data set. Each heat map illustrates the transformation of the true empirical correlations *corr* (*Y*_*n*_, *Y*_*m*_) (horizontal axis) after adding bias and applying the corresponding normalization method. In the top left plot the vertical axis represents the observed correlations 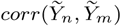. In the remaining five heat maps, the vertical coordinates represent the correlations after normalization. Going clockwise, these five heat maps are Rank, Quantile, MAD, LOESS, and C-SHIFT. The darker the color, the higher the density. The number on top of each heat map indicates the relative leftover error after normalization. Smaller numbers indicate better recovery performance.

In Figures 1 and 3 we depict the bar graphs of correlations for RCM and Cascade data sets, respectively. On the x-axis we display the range of correlations, partitioned into the intervals of length 0.1. The height of each bar represents the number of correlations that belong to the corresponding interval. Bars of different colors correspond to different correlation matrices, indicated in the legend. In particular, black and yellow bars correspond to the correlation matrices of empirical and observed data, respectively. As we can see in both data sets, the correlations of the observed data (yellow) are shifted away from −1 so that there are no large magnitude negative correlations. The aim of the normalization algorithms is to shift the correlations back into correct positions, i.e., ideally, the correlations of the normalized data should match the empirical correlations. Note that for both data sets, the C-SHIFT method correctly recovers the number of correlations in each interval: the red bars almost perfectly match the black bars. In contrast, other normalization methods could not recover the correct numbers of correlations, especially for the correlations of larger magnitudes. Specifically, Rank, Quantile and LOESS normalization techniques tend to shift correlations mostly to the center of the bar plot, each forming a bell shape. Predictably, the MAD method has the worst performance in correlation recovery. Finally, among the other three normalization techniques (Quantile, Rank, and LOESS), the latter method has the poorest performance.

Figures 2 and 4 contain six heat maps each, for RCM and Cascade data sets, respectively. Each heat map illustrates the transformation of the true empirical correlations *corr*(*Y*_*n*_, *Y*_*m*_) (horizontal axis) after adding bias and applying the corresponding normalization method. We consider 2,001,000 correlations corresponding to all pairs of genes. For each point, representing a pair of genes (*n, m*), the horizontal coordinate equals the true empirical correlation *corr*(*Y*_*n*_, *Y*_*m*_) in all six plots. The vertical coordinate in the top left heat map is the correlation in the observed data, 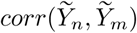. Importantly, it shows the shift of correlations rightward in the observed data. In the remaining five heat maps, the vertical coordinates represent the correlations after normalization. Going clockwise, these five heat maps are Rank, Quantile, MAD, LOESS, and C-SHIFT. The darker the color, the higher the density. Notice that the heat map for C-SHIFT is almost perfectly diagonal, which demonstrates how well C-SHIFT recovers the correlations. Thus, in addition to correctly recovering the right numbers of correlations in each interval (which was demonstrated in Figures 1 and 3), the proposed C-SHIFT algorithm also returns (shifts back) the correlations to the correct margins. Hence, the heat map is a diagonal line. The number on top of each heat map indicates the relative leftover error after normalization, i.e., the *ℓ*^2^-norm of the vector of differences between the horizontal and vertical coordinates, scaled by the Frobenius norm of the difference between the empirical and the observed correlation matrices. Thus, the left top heat map is assigned the value 1, and for each normalization method, the smaller the number the better it recovers the empirical correlation matrix. Any such number smaller than one is an improvement. The number for C-SHIFT is by far the smallest in each data set (0.023518 and 0.023881), while in the case of MAD normalization, the corresponding number even exceeds 1.

## III. Discussion

In systems biology, the gene co-expression networks (GCN) are reconstructed from the correlations between the genes. GCN recovery relies on removing the bias with a normalization method, and thus improving the estimation of correlations between the pairs of genes. However, the standard normalization techniques such as Rank, Quantile, LOESS, and MAD are known to be insufficient at recovering the true empirical correlations while the C-SHIFT algorithm is specifically designed to recover the true empirical correlations. The multiple experiments with synthetic data sets demonstrate the algorithm’s superior performance in comparison to the standard normalization techniques.

Importantly, we notice that the C-SHIFT algorithm corrects the positive shift of covariances (and correlations) observed when 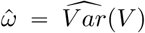 is larger than 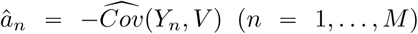 in (7). Hence, the independence of *V* from *Y*_*n*_ assumption can be replaced with a weaker assumption stating that *Cov*(*Y*_*n*_, *V*) ≪ *Var*(*V*). This will be explored in a follow-up publication.

An alternative version of the C-SHIFT algorithm is based on trace minimization approach instead of energy minimization. In this alternative C-SHIFT algorithm, the positive semi-definite matrix 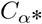 with

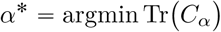

is used to approximate the true empirical covariance matrix *C*. The analogs of Lemmas 1 and 2 and the convexity result in Theorem 1 are also established for Tr(*C*_*α*_) in the trace minimization approach. See [11]. Empirically it appears that this alternative approach produces the same *α*^*^ as the original C-SHIFT algorithm based on energy minimization as presented in this paper, and therefore it recovers the empirical covariance *C* with the same accuracy. Thus, the alternative, trace minimizing C-SHIFT algorithm can be used instead of Algorithm 1. This approach will be analyzed in a follow-up paper.

Finally, the C-SHIFT algorithm was deposited on GitHub at https://github.com/prlogan/C-SHIFT

## Acknowledgments

This research is supported by the FAPESP awards 2018/14952-7 and 2018/07826-5, and by the NSF award DMS-1412557.

## Footnotes

* In this paper we will refer to the phenomenon as the blessing of dimensionality rather than the curse of dimensionality.

